# Patient-Derived Medullary Thyroid Cancer Organoids; a Model for Patient-tailored Drug and PET-Tracer Screening

**DOI:** 10.1101/2023.09.18.558266

**Authors:** Luc H.J. Sondorp, Eline C. Jager, Inês F. Antunes, Rufina Maturi, Liesbeth Jansen, Wouter T. Zandee, Adrienne H. Brouwers, Thera P. Links, Robert P. Coppes, Schelto Kruijff

**Affiliations:** Department of Surgical Oncology, University of Groningen, University Medical Center Groningen, Groningen, The Netherlands; Department of Biomedical Sciences of Cells and Systems, Section Molecular Cell Biology, University of Groningen, University Medical Center Groningen, Groningen, The Netherlands; Department of Endocrinology, University of Groningen, University Medical Center Groningen, Groningen, The Netherlands; Department of Nuclear Medicine and Molecular Imaging, University of Groningen. University Medical Center Groningen, Groningen, The Netherlands; Department of Molecular Medicine and Medical Biotechnology, University of Naples Federico II, Naples, Italy; Department of Radiation Oncology, University of Groningen, University Medical Center Groningen, Groningen, The Netherlands; Department of Molecular Medicine and Surgery, Karolinska Institutet, Stockholm, Sweden

## Abstract

**Background:** Medullary thyroid carcinoma (MTC) is a neuroendocrine tumor derived from the parafollicular C-cells of the thyroid gland. PET imaging, with various PET tracers, is performed when distant metastatic disease is suspected. After the recognition of progressive disease on imaging, targeted therapy may be initiated to prolong survival. Mutations in the gene encoding the REarranged during Transfection (RET) tyrosine kinase play a key role in the development of MTC. It seems that tyrosine kinase inhibitors (TKIs) inhibit tumor proliferation, but it remains challenging to determine the best patient specific treatment option. Here, we aim to set up an *in vitro* MTC organoid model to study its potential for patient-tailored drug-screening and uptake of PET tracers.

**Methods:** Dispersed cells obtained from surgical MTC biopsies were suspended in Matrigel with defined medium allowing MTC organoid formation. To study putative MTC stem cells, the self-renewal potential of organoids was tested by dissociation to single cells and re-plating. To check MTC origin, MTC-specific gene expression and proteins were characterized by qPCR and immunofluorescent (IF) staining. To investigate cytotoxicity, MTC-organoids (MTOs) were exposed to various TKIs after which hormone (calcitonin and CEA) excretion levels were determined. Lastly, we evaluated cell-specific uptake of clinically used Positron Emission Tomography (PET) tracers.

**Results:** Nine MTC biopsies were processed and cultured as MTOs. Eight MTO lines were used to determine organoid formation efficiency (OFE), which yielded a maximum OFE of 6.3% in passage 1 (p1), 5.9% in p2, and 9.4% in p3, indicating the presence of putative stem cells. IF staining showed expression of MTC-specific markers in both tissue and MTOs showing tissue resemblance. Tumor marker measurements in MTO medium showed MTC-specific production of calcitonin and CEA with changed concentrations after exposure to TKIs. Exposure to PET tracers showed significant uptake in the MTOs.

**Conclusion:** MTC organoids can be successfully cultured and resemble the tissue of origin in gene expression, protein expression and functionality. In addition, MTOs can take up PET tracers, and have the potential to be used as a prediction model for TKI treatment in the future.

## Introduction

Medullary thyroid cancer (MTC) is a rare type of thyroid cancer, derived from the parafollicular C-cells that lie between the thyroid hormone producing follicles of the thyroid gland (1). MTC is a neuro-endocrine tumor that constitutes 5% of thyroid cancers and occurs sporadically (65-75%) or hereditarily (25-35%) (2–4).

Genetic mutations in *REarranged during Transfection* (*RET*) play a vital role in the development of all hereditary MTCs and 65-100% of sporadically occurring MTCs (5–8). *RET* encodes a tyrosine kinase that activates numerous intracellular proteins through phosphorylation. However, mutations in *RET* cause phosphorylation of the tyrosine kinase with enhanced signal transduction of downstream pathways PI3K-AKT and RAS-MAPK and cancer cell proliferation (9,10).

Once the disease has become clinically apparent, most patients undergo direct surgical treatment. In a curative setting, total thyroidectomy combined with lymph node dissection of the central compartment (level VI) are standard of care. Worldwide, additional dissections of the lateral neck compartment(s) (level 2-5b) are performed when preoperative tumor markers (calcitonin and carcinoembryonic antigen [CEA]) are disproportionally high and/or metastasized lymph nodes are detected on preoperative imaging (11).

Advances in molecular imaging techniques have improved pre- and postoperative tumor staging in MTC patients. ^18^F-fluorodeoxyglucose (^18^F-FDG) Positron Emission Tomography (PET) and ^18^F-dihydroxyphenylalanine (^18^F-DOPA) PET are used in clinical practice to assess the extent of disease. Where ^18^F-FDG has an overall sensitivity of 62-76% in MTC patients (12,13), ^8^F-DOPA PET has a slightly higher overall sensitivity of 44-83% for MTC and allows better detection of disease in patients with indolently growing MTC(14). However, neither PET tracer detects all MTC lesions in all patients reliably, which makes it difficult to determine the best PET tracer for each individual patient making pre-clinical models to screen, test, and validate (new) tracers an unmet need.

Furthermore, when distant metastatic disease is apparent, systemic therapy may be considered to prolong progression-free survival. There are currently four systemic options for patients with MTC; all are tyrosine kinase inhibitors (TKIs) that inhibit intracellular signal transduction pathways by blocking tyrosine kinases (15)(9) The latter TKIs have shown promising results in phase I and II clinical trials(16,17). Although the development of new TKIs has enlarged the treatment reservoir, it has also brought about the challenging to determine individual patient’s most adequate therapy.

A patient-derived organoid model may guide patient-specific imaging and treatment. Cancer stem cells are believed to be a subpopulation within malignant tumors responsible for cell growth and dedifferentiation (18). After a tissue biopsy, these adult cancer patient specific stem cells can be obtained and cultured to form *in vitro*, 3-dimensional spheres. These patient-derived spheres, also known as patient-derived organoids, mimic the in vivo situation in a realistic way and can be used to study metastatic potential, genetic alterations, tumor heterogeneity and treatment sensitivity (19–21). This study therefore aims to culture MTC organoids (MTOs) for the first time and evaluate their potential for treatment sensitivity testing and for PET tracer development.

## Results

### Patient characteristics

Nine patients with MTC at diagnosis were included into the study after giving informed consent. The median age at time of inclusion was 64 (36 – 69) years, and three patients were female. In all patients, lymph node metastases were found. Out of all these, three patients presented with additional distant metastases. Germline *RET* analysis was performed in eight of the patients, which established the hereditary Multiple Endocrine Neoplasia type 2A (MEN2A) syndrome in two patients. In one patient, no germline RET analysis was available, and the disease was classified as sporadic MTC, based on a negative family history and absence of other MEN2 related diseases. In one patient with sporadic disease, the presence of a somatic *RET* mutation was assessed to evaluate eligibility for a clinical trial. All patients were surgically treated, two (22%) patients received a TKI additionally, and in 2 (22%) patients, postoperative radiotherapy was applied.

**Table 1.** Characteristics of patients included in this study. Abbreviations; MTC = medullary thyroid cancer; RET = rearranged during transfection (proto-oncogene); MEN2A/B = Multiple Endocrine Neoplasia type 2A/B.

| Patient characteristics | MTC patients<br>n (%) or median (IQR) |  |
| --- | --- | --- |
| **Sex** |  |  |
| Female | 3 | (33) |
| Male | 6 | (66) |
| **Age at time of inclusion (years)** | 64 (36 – 69) |  |
| **Status at last follow-up** |  |  |
| Alive | 7 | (78) |
| Deceased | 2 | (22) |
| **Extent of disease at time of inclusion** |  |  |
| Lymph node metastases | 9 | (100) |
| Distant metastases | 3 | (33) |
| **Type of MTC** |  |  |
| Sporadic disease | 7 | (77) |
| MEN2A | 2 | (22) |
| **Genetic analyses** |  |  |
| Germline *RET* mutation | 8 | (89) |
| Cys634Arg mutation | 2 | (25) |
| No germline mutation present | 5 | (63) |
| Somatic *RET* mutation | 1 | (11)* |
| Met918Thr mutation | 1 | (100) |
| **Treatment Modalities** |  |  |
| Surgery | 9 | (100) |
| Tyrosine Kinase Inhibitor | 2 | (22) |
| Radiotherapy | 2 | (22) |

\*Analysis of somatic *RET* mutations was not standard of care in clinical practice and only performed when there was a clinical indication (in one patient only).

### Organoid formation and in vitro self-renewal potential of medullary thyroid carcinoma stem cells

Nine medullary thyroid carcinoma (MTC) biopsies were dissociated into dispersed cells using mechanical and enzymatic digestion. These cells were cultured in Matrigel with defined medium, and formed spheroid structures (**Figures 1A and 1B**, passage 0). To determine the presence of potential stem cells, self-renewal potential of single cells was assessed. For this, cultures from eight patients were used to determine Organoid Formation Efficiency (OFE), with all lines reaching passage 1 (p1), 87.5% reaching p2, and 62.5% reaching p3. A maximum OFE of 6.3 ± 0.81% in p1, 6.9 ± 0.83% in p2, and 9.4 ± 1.17% in p3 was reached (**Figures 1B and 1C**), showing self-renewal potential of medullary thyroid carcinoma cells. These spheroids continued to secrete calcitonin and CEA in all passages indicative of MTC derived organoids (MTO), although a decline in concentration was seen from p1 to p2 and p3. This is possibly due to a gradual decrease in the number of differentiated cells, with only stem cells remaining. (**Figure 1D**).

**Figure 1.**
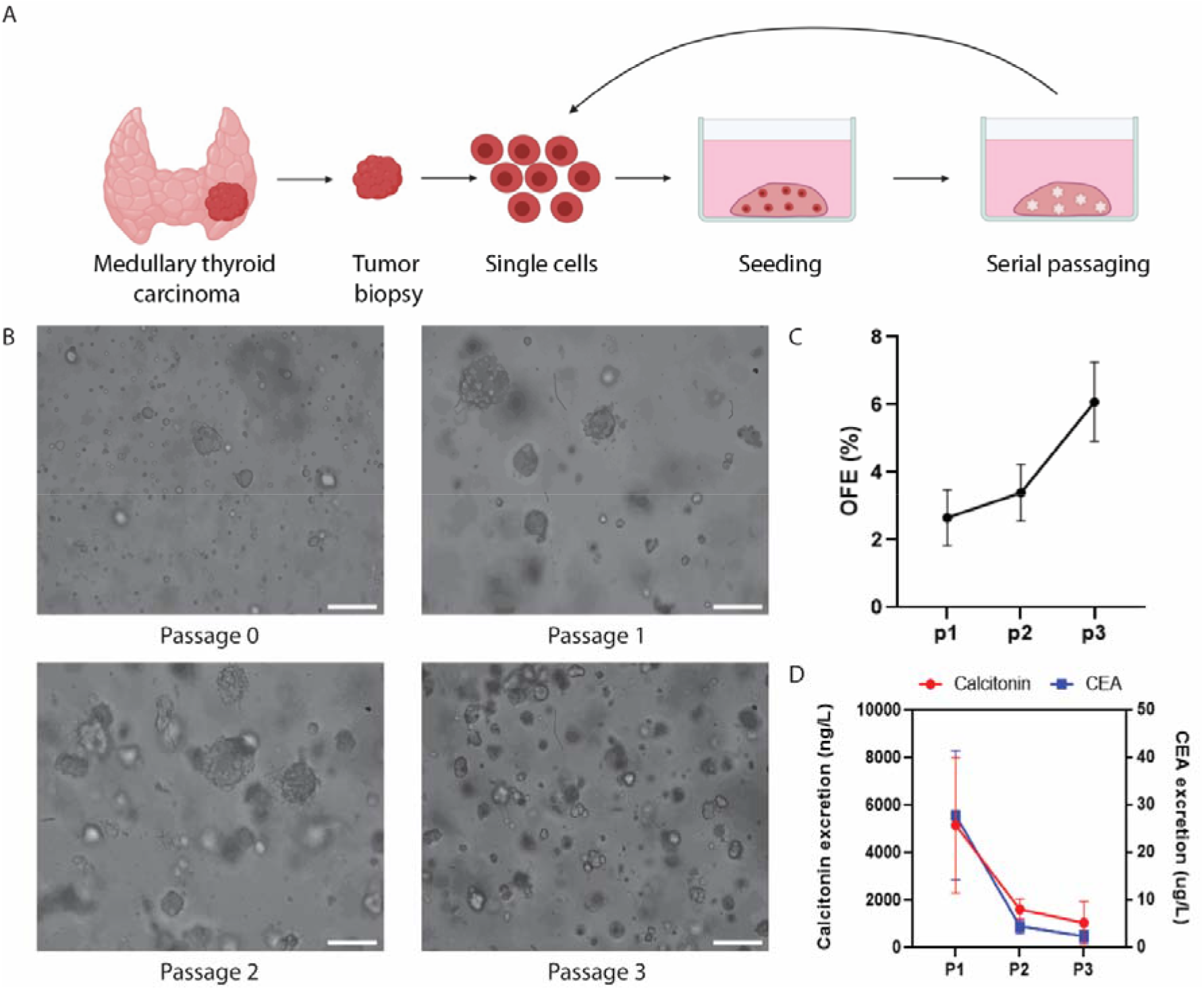
In vitro self-renewal of organoid-forming cells from human putative medullary thyroid cancer stem cells. (A) Schematic of medullary thyroid cancer tissue isolation, primary tissue culture, and the self-renewal assay. (B) Primary MTC-thyrospheres 7 days in culture (p0) and MTOs at the end of p1, p2, and p3. Scale bars, 100 μm. (C) Self-renewal potential of MTOs during passaging (n = 9 patients; error bars represent range). (D) In medium secreted calcitonin and CEA levels from MTOs during passaging, calcitonin is plotted on the left y-axis, and CEA on the right y-axis (n = 3 patients; error bars represent SEM).

### Characterization of MTOs

To further characterize the MTOs, we selected four MTC specific genes: *Calcitonin Related Polypeptide Alpha (CALCA), Carcinoembryonic Antigen-related Cell Adhesion Molecule (CEACAM), Thyroid Transcription Factor 1 (TTF1)*, and *Somatostatin (SST)*. qPCR analysis showed that all genes were found in the originating tissue and derived MTOs, although with reduced expression levels in MTOs, as can be expected from less differentiated cells **(Figure 2A)**. This is in concordance with other (para)thyroid derived organoid models developed by our group (20,22). Immunofluorescent labeling and imaging confirmed corresponding MTC-specific protein expression in tissue and MTOs **(Figure 2B)**. CALCA and CEACAM encode the tumor-markers which are known to be secretary products of MTCs. When comparing the expression patterns of CALCA and CEACAM in tissue and MTOs, a widely dispersed expression pattern is seen in tissue, whereas the MTOs show intensive expression along the outer edges for CALCA and confluent cytoplasmic expression of CEACAM. TTF1 is a nuclear marker in thyroid originating tissue and is crucial for thyroid differentiation and morphogenesis (23). TTF1 immunolabeling shows nuclear expression in tissue, indicative of an active proliferative state, but a mostly cytoplasmic expression pattern in the MTOs, being the inactive state. Somatostatin is another excreted factor that is present in MTC tissue and MTOs, which results in the widely dispersed pattern observed in both tissue and MTOs.

**Figure 2.**
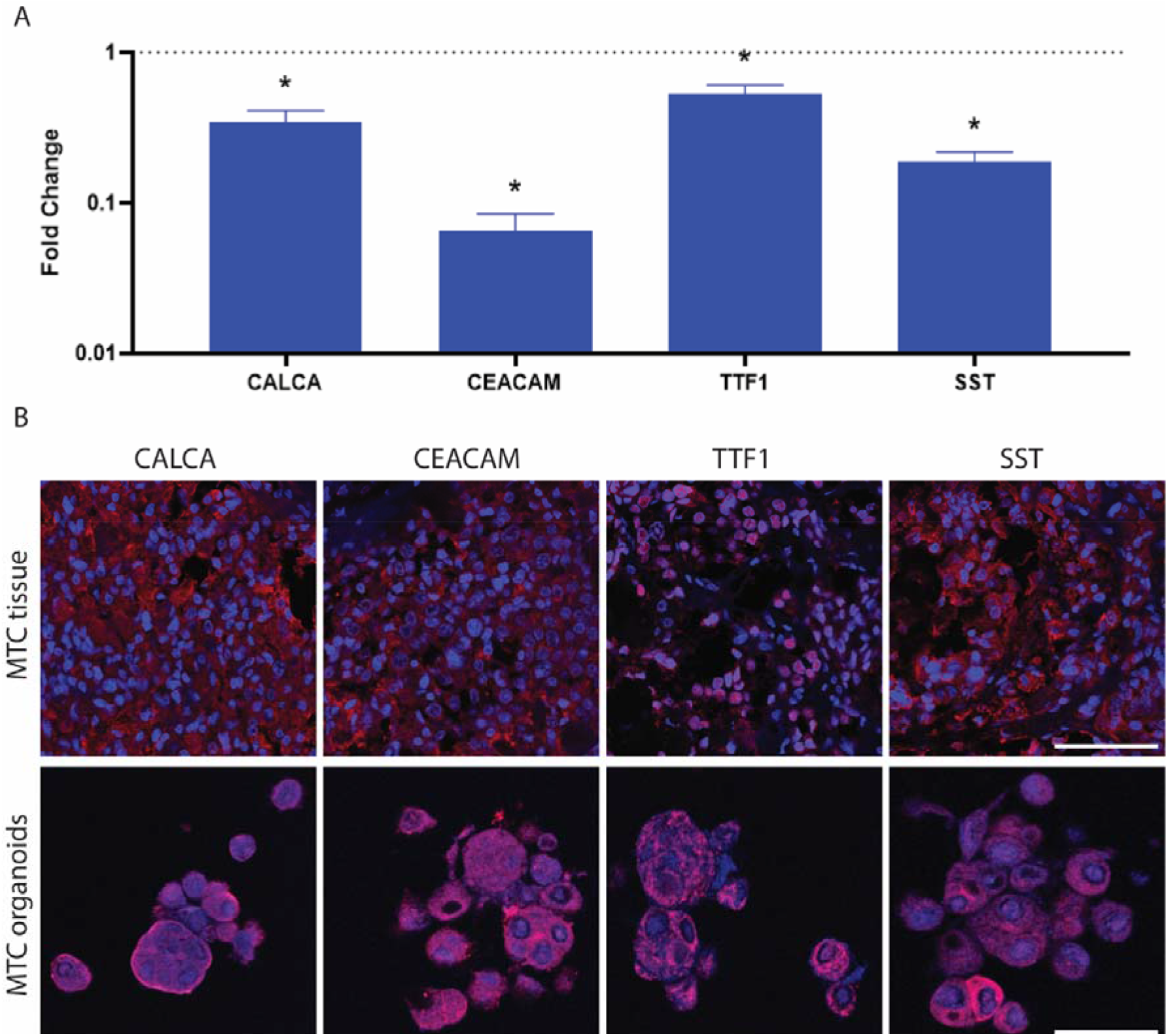
Immunofluorescence characterization of MTOs. A. qPCR analysis of the MTC specific markers in MTOs. B. Representative images of organoids and tissue, showing MTC-specific markers. Scale bars, 70 μm (tissue) and 20 μm (organoids). Markers indicated at the top are shown as a red fluorescent signal. Nuclei are shown as a blue fluorescent signal. (error bars represent the SEM of three biological replicates, normalized to MTC tissue = dotted line. * p < 0.05).

### Drug Screening

To evaluate the potential of patient-derived MTOs in patient-tailored TKI treatment selection, TKIs were administered to assess the effect on the secretion of tumor markers, calcitonin and CEA (**Figure 3A**).

**Figure 3.**
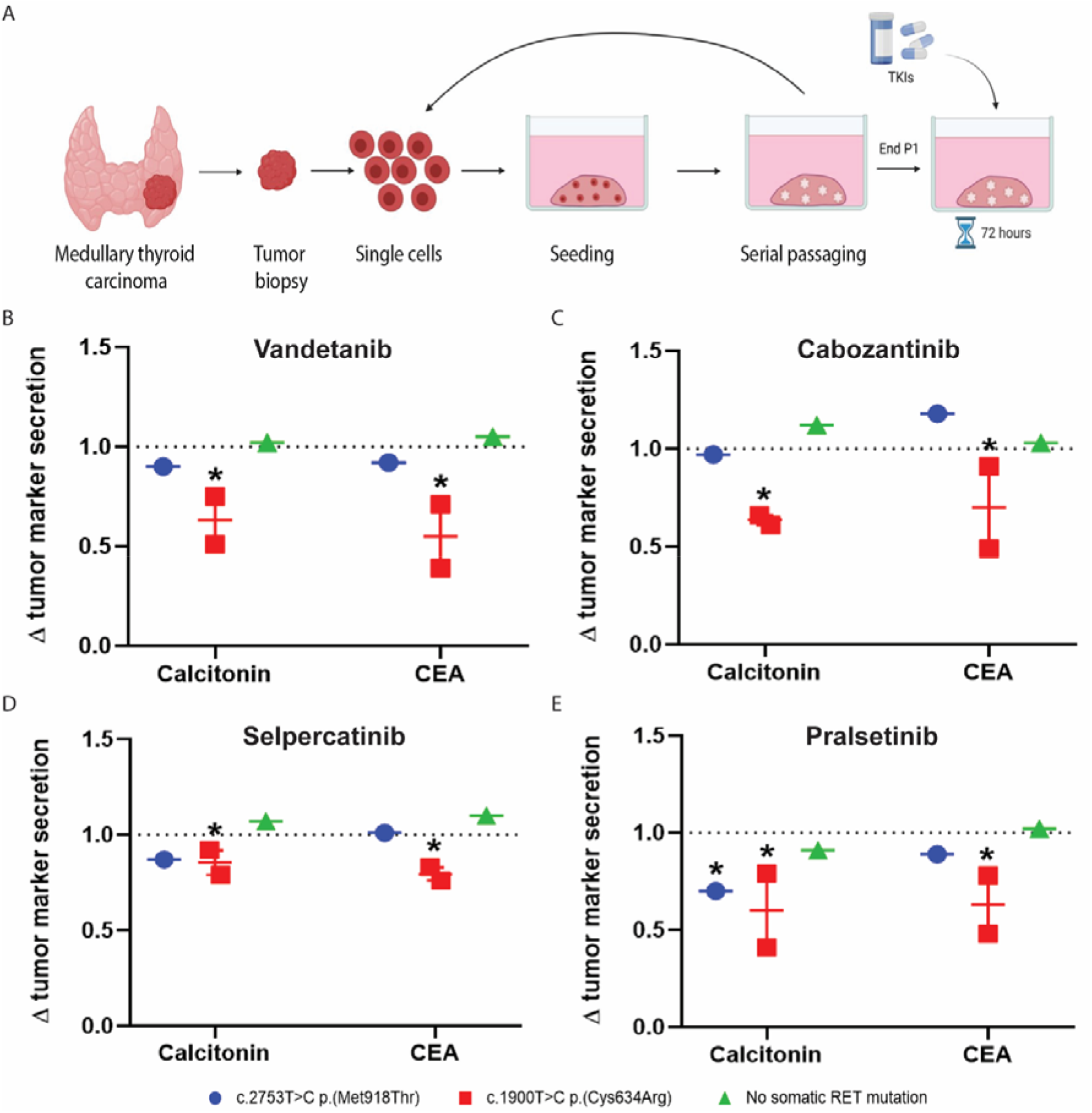
Effect of TKIs on Calcitonin and CEA concentrations in culture medium from MTOs. (A) Schematic of MTC tissue isolation, primary tissue culture, and the drug screening experiment. MTOs exposed to: (B) Vandetanib. (C) Cabozantinib. (D) Selpercatinib. (E) Pralsetinib. All; n = 4 patients; error bars represent SEM, normalized to DMSO controls, dotted lines * p < 0.05.

In order to select the appropriate dose for this proof-of-concept drug screening, spheroids were cultured using the MTC cell line MZ-CRC-1. These spheroids were treated with a logarithmic dose de-escalation of vandetanib, cabozantinib, selpercatinib, and pralsetinib. The Incucyte live-cell analysis system was used to determine the IC50 value of each TKI (**Supplemental Figure 1**). The final dose of 1.0 μM of vandetanib and 0.1 μM of cabozantinib, selpercatinib, and pralsetinib were added to the medium of MTC organoids at the end of passage 1. After 72 hours, medium was collected to measure concentration of secreted calcitonin and CEA.

We observed a significantly decreased calcitonin concentration in the medium of organoids cultured from the two patients with a Cys634Arg RET mutation, when exposed to either of the four TKIs tested and compared with the DMSO control (average reduction of 37 ± 8.6% when exposed to vandetanib, a 36.5 ± 9.1% reduction when exposed to cabozantinib, 14.5 ± 3.1% when exposed to selpercatinib and 40 ± 10.5% when exposed to pralsetinib) (**Figure 3B-E**). A similar effect was observed in the secretion of CEA in the same MTOs (average reduction of 45 ± 8.6% when exposed to vandetanib, a 30 ± 9.2% reduction when exposed to cabozantinib, 20.5 ± 3.2% when exposed to selpercatinib and 37 ± 10.5% when exposed to pralsetinib). The organoids obtained from the patient with the somatic Met918Thr RET mutation showed a significant reduction (30 ± 13.5%) in calcitonin secretion when exposed to pralsetinib (**Figure 3E**).

Our results suggest a possible mutation-specific response of our patient-derived organoids to the four TKIs that were used. Within the group of patients with a Cys634Arg mutation (2 patients), we noticed a heterogeneous change in tumor marker secretion levels. This might suggest that other factors, in addition to the specific RET mutation, play a role in TKI response. Altogether, these data suggest a potential of MTOs for patient-tailored drug screening.

### Positron Emission Tomography (PET) Tracer Validation

To demonstrate the utility of patient-derived MTOs to validate PET tracers, ^18^F-FDG, ^18^F-DOPA, and ^18^F-PSMA, were used to assess uptake by MTOs (**Figure 4A**). In contrast to ^18^F-FDG and ^18^F-DOPA, ^18^F-PSMA PET has no role in MTC detection in clinical practice. However, a few case reports have illustrated the detection of MTC and ^68^Ga-PSMA PET scanning which led to our interest in this tracer (24–26). Moreover, in our center, we use ^18^F-PSMA instead of ^68^Ga-PSMA. Both^18^F-PSMA and ^68^Ga-PSMA target the prostate-specific membrane antigen (PSMA), which is a transmembrane protein, and therefore a useful diagnostic and, possibly, therapeutic target.

**Figure 4.**
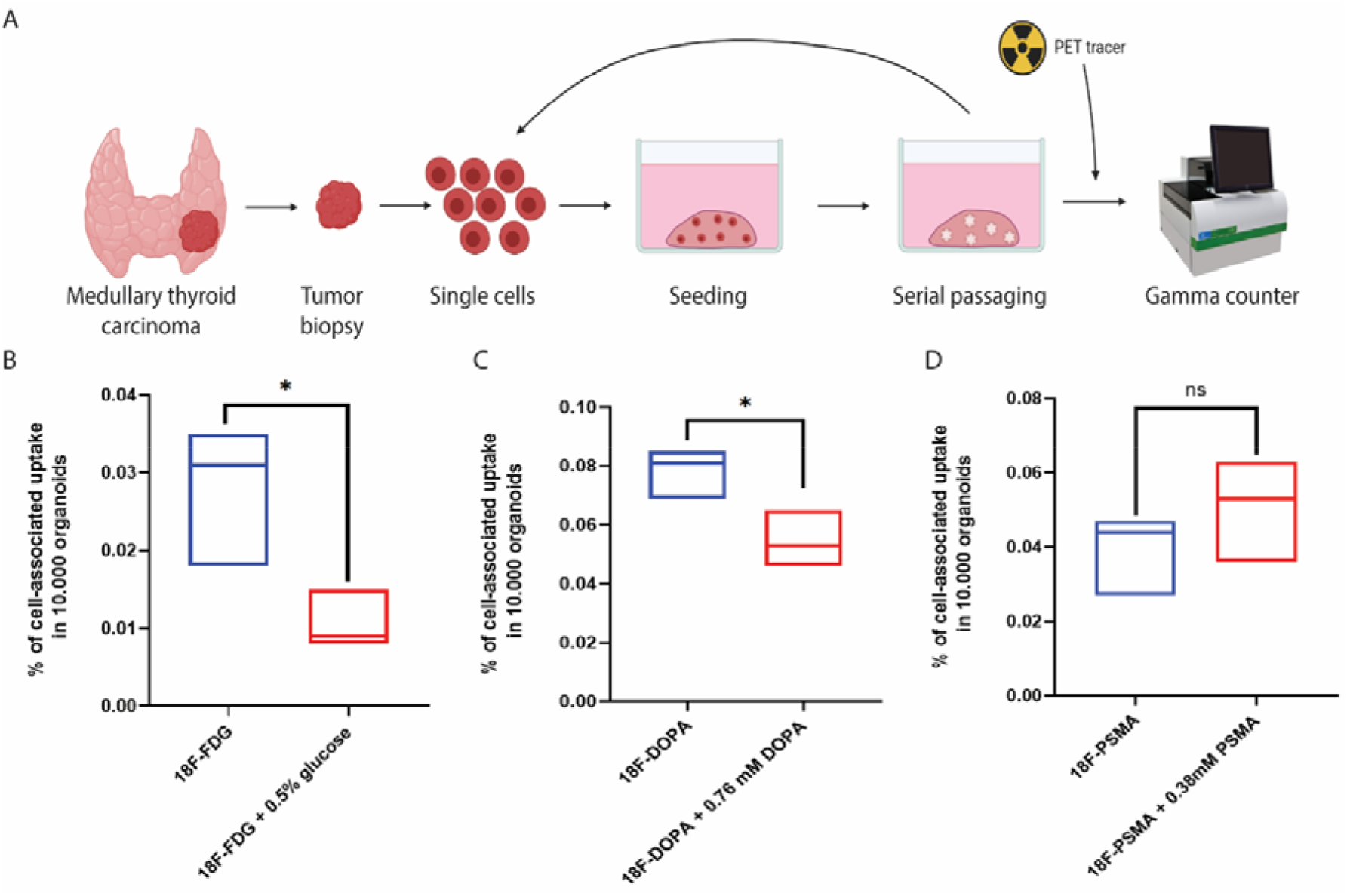
PET experiment of MTOs with radioactive tracers. (A) Schematics of MTC tissue isolation, primary tissue culture, and the PET/SPECT experiment. (B) Percentage of ^18^F-FDG cell-associated uptake compared with the percentage of cell-associated uptake after saturation using 0.5% glucose (n = 3 patients, Tukey boxplot). (C) Percentage of ^18^F-DOPA cell-associated uptake compared with the percentage of cell-associated uptake after saturation using cold DOPA (n = 3 patients, Tukey boxplot). (D) Percentage of ^18^F-PSMA cell-associated uptake compared with the percentage of cell-associated uptake after saturation using cold PSMA (n = 3 patients, Tukey boxplot).

We found that the MTOs showed cell-associated uptake of ^18^F-FDG (cell-associated uptake in 10,000 organoids, 0.028 ± 0.009%), ^18^F-DOPA (cell-associated uptake in 10,000 organoids, 0.078 ± 0.008%), and ^18^F-PSMA (cellular uptake in 10,000 organoids, 0.039 ± 0.011%) (**Figures 4B-D**). ^18^F-FDG showed a significantly reduced cell-associated uptake when MTOs were first saturated with 0.5% glucose (0.028 ± 0.009% versus 0.011 ± 0.004%), indicating specific uptake (**Figure 4B**). Similarly, a significant reduction in the uptake of ^18^F-DOPA was observed when MTOs were saturated with an excess of cold DOPA (0.078 ± 0.008% versus 0.055 ± 0.010%, p = 0.032). This also indicates specific uptake of ^18^F-DOPA (**Figure 4C**). However, when we saturated the uptake of ^18^F-PSMA, with an excess of cold PSMA, we observed no significant difference (0.039 ± 0.011% versus 0.051 ± 0.014%).

Our results indicate cellular uptake of ^18^F-FDG and ^18^F-DOPA by the MTOs. Accumulation of ^18^F-FDG in the MTOs suggests a higher glucose metabolism, which is commonly found in tumor cells (27). ^18^F-DOPA binds to the LAT-1 transporter (amino acid transporter) commonly found on neuroendocrine cells, allowing internalization of the tracer (28). MTC is derived from neural crest cells and therefore of endocrine origin, which could clarify specific uptake of the MTOs. On the contrary, ^18^F-PSMA binds to PSMA which is only found on the neo-vasculature of MTCs (29). Absence of vascularization in MTOs could therefore explain non-specific uptake of ^18^F-PSMA. Altogether our results show potential of MTOs to understand PET tracer uptake mechanisms. Moreover, MTOs may be used in PET tracer development.

## Discussion

MTC is a rare type of thyroid cancer derived from the parafollicular C-cells, mainly treated with surgery and/or systemic therapy with TKIs. It has been shown that a patient-specific approach is fundamental to prolong survival (30). In the present study, we show the development and self-renewing capacity of patient-derived medullary thyroid cancer organoids (MTOs) and illustrate potential applications after culturing organoids from this rare endocrine tumor. Characterization of MTOs and their originating tissues has revealed similarities in terms of MTC-specific markers on both gene and protein level. In addition, we note that MTOs mimic the originating tissue in functionality by producing specific tumor markers: calcitonin and CEA. Moreover, our results show specific uptake of PET tracers by the MTOs, and a response to TKIs in the MTOs of patients with a Cys364Arg RET mutation. Altogether our results show representative and functional patient-derived MTC organoids.

For development and testing of new imaging tracers and drugs, we aimed to establish an appropriate pre-clinical model for testing. The model had to be robust, reliable, quick in use, and able to allow high-throughput screening methods. When we started to work on the MTOs and exposed them to clinically available TKIs, we observed a heterogeneous response, demonstrating that these organoids can mimic MTC functionality and are a valid in vitro model for drug testing. We also demonstrated that the MTOs revealed specific uptake of the PET tracers ^18^F-FDG and ^18^F-DOPA that are currently being used to evaluate MTC patients. This further validates their origin from MTC cells and indicates that we may be able to culture patient-specific MTOs, to use them in the future as a drug and PET tracer testing platform. The development of such methodology could ultimately allow patient response prediction using patient-derived MTC organoids. By assessing response to systemic therapy in MTOs prior to patient treatment initiation, our aim is eventually to prevent ineffective treatments and unnecessary adverse events. Moreover, MTO cultures could guide the development and testing of effective MTC imaging tracers. By determining the PET tracer with highest uptake in a patient-derived preclinical model, PET imaging may become personalized, lead to improved disease staging at diagnosis and in the follow-up, and treatment may be adapted accordingly.

This study is the first to develop a patient-derived MTO culture methodology of medullary thyroid cancer tissue. Organoids are self-organizing 3D culture systems that are highly similar to actual human organs, including increased exposure to circulating molecules, neighboring cells, and the extracellular matrix (Matrigel) (31,32). Organoids from several endocrine organs and endocrine cancers have been developed, such as thyroid organoids (33,34), papillary thyroid cancer organoids (20), and parathyroid organoids (22). Recently, numerous papers have shown the value of organoids for discovering potential novel therapeutic targets, at the same time also opening opportunities for future imaging tracer testing and targeted treatment screening (19,22).

The limitations of our study include absence of the original microenvironment. This consists of, among others, fluctuating concentrations of extracellular signals and the presence of blood vessels. This may be the reason why our MTOs did not reveal ^18^F-PSMA uptake. Several case reports showed uptake of PSMA-binding tracers in MTC which led to our experiment. However, histological examination of MTC tissue shows PSMA expression only in the neo-vasculature of the tumor, which are not incorporated in the organoids and may therefore explain the absence of 18F-PSMA uptake in our organoids (29). Although a more complex organoid model may improve the resemblance to in vivo tissue, bioengineers have suggested that a robust model, with the specific characteristics relevant for the (patho)physiological functions of interest *only*, may be sufficient for the study of mechanisms and functions (35).

Subsequently, when MTOs were exposed to TKIs, we observed a significant decrease of biomarkers in patients with a Cys634Arg mutation. It is difficult to directly compare the results of these experiments to patients. In contrast to patients, who take daily oral doses of a TKI, we chose to administer only a single dose of each TKI to the MTOs. In addition, we measured the tumor marker concentrations after 72 hours, according to our earlier patient-derived organoid drug screening experiments, purely to show a proof of principle mechanism (22). Moreover, it is difficult to extrapolate the administered dose to a patient dose. Pharmacokinetic characteristics, possibly drug-drug interactions and renal-or hepatic impairment in patients influence how much of the administered TKI dose finally reaches the tumor cells. Additionally, some suggest that decreases of calcitonin and CEA do not adequately reflect response to TKIs (36)(36). Measurements on CT imaging are therefore the gold-standard for assessing tumor response in clinical practice. However, growth rate is extremely difficult to determine in organoids. One passage taking three weeks and no growth rate could therefore be determined within the 72 hours of TKI treatment. However, measuring differences in ATP levels could provide more information about the metabolic activity of the TKI exposed organoids in future studies.

In conclusion, we present a patient-derived MTO culture that recapitulates the originating tissue in terms of gene and protein expression and functionality. This medullary thyroid cancer culture may be able to serve as a novel in vitro model to perform physiology studies and discover potential novel therapeutic targets, opening opportunities for future tracer testing and drug screening.

## Materials & Methods

### Patient material

Human MTC tissue was obtained from patients undergoing surgical treatment. The biopsies were transported from the operating room (OR) in Hank’s Balanced Salt Solution (HBSS) (Gibco, NY, USA) with 1% bovine serum albumin (BSA) (Gibco), on ice for immediate further processing.

### MTC-thyrosphere cultures

The biopsies were mechanically dispersed using the gentleMACS dissociator (Miltenyi Biotec, Leiden, the Netherlands) and simultaneously subjected to digestion in 5mL HBSS/1% BSA buffer containing 80 mg/mL dispase (Sigma), 1.2mg/mL collagenase type II (Gibco), and calcium chloride (Sigma) at a final concentration of 6.25□mM. This was followed by two periods of 15 minutes in a 37°C shaking water bath. Hereafter, the isolate was collected by centrifugation, washed in HBSS/1% BSA solution, and passed through a 100□μm cell strainer (BD Biosciences, NJ, USA). The cell pellet was collected by centrifugation and resuspended in Dulbecco’s modified Eagle’s medium: F12 medium (DMEM:F12) containing Pen/Strep antibiotics (Invitrogen) and Glutamax (Invitrogen). Every 20 μL of this cell solution was combined with 40 μL of Basement Membrane Matrigel (BD Biosciences) on ice and subsequently plated in the center of a 24-well tissue culture plate. Following solidification of the gel for 30 minutes in the incubator, 500 μL of complete growth medium (MTC-medium) consisting of 50% conditioned Wnt3a medium, 10% conditioned R-Spondin medium and 40% DMEM:F12 containing B27 (Gibco, 0.5x), HEPES (Gibo, 10 mM), ROCK inhibitor Y-27632 (10 μM, Abcam, Cambridge, UK), Nicotinamide (100 uM, Sigma), Noggin (25 ng/mL, Peprotech, NJ, USA), EGF (20 ng/mL, Sigma), FGF-2 (20 ng/mL, Peprotech), TGF-β inhibitor A 83-01 (5 uM, Tocris bioscience), Fungin (10μg/ml, Invivogen), and VEGF-121 (100 ng/mL, Immunotools) was added to each well and replaced on a weekly basis.

### Self-renewal

To study the long-term self-renewing potential, organoids were dissociated and re-plated for the next passage. Prior to passaging, Matrigel was dissolved by incubation with dispase enzyme (1□mg/mL for 30 minutes at 37 °C; Sigma) and organoids with a size >50 μm were counted using the Ocular Micrometer at a 10x objective. Initial MTC-thyrosphere cultures of seven days old (d) were dissociated to single cells using 0.05% trypsin-EDTA (Invitrogen), counted using Tryphan blue and cell concentration was adjusted to 2 x 10^6^ cells per ml. In total, for every 20 μL of this cell solution, 40 μL of Basement Membrane Matrigel (BD Biosciences) was added while on ice, and plated in the center of a 24-well tissue culture plate. After solidifying the Matrigel for 30 minutes at 37 °C, gels were covered in 500 μL of medium as defined above and this medium was renewed weekly. This self-renewal procedure was repeated every three weeks and up to 4 times (4 passages).

### Cell Culture

The human medullary thyroid cancer cell line MZ-CRC-1, was cultured in DMEM medium supplemented with 10% fetal calf serum (Thermo Scientific, Paisley, UK) and Pen/Strep (Invitrogen, Bleiswijk, The Netherlands).

### Calcitonin and CEA secretion

Medium was collected before passage and after TKI treatment and stored at -80 °C until use. Medium was used to measure the concentration of secreted calcitonin and CEA protein by the MTOs. Secreted calcitonin was measured by electrochemical luminescence immunoassay (ECLIA) using an Elecsys Calcitonin kit (Roche, Germany). Secreted CEA was measured by chemiluminescent microparticle immunoassay (CMIA) using an Alinity CEA kit (Abbott Ireland Diagnostics).

### Quantitative Polymerase Chain Reaction

Total RNA from patient material (n=3) and organoids (xxx) was extracted (RNeasy™ Mini Kit, Qiagen). To obtain cDNA, 500ng of total RNA was reverse transcribed using 1 μL 10 mM dNTP Mix, 100 ng random primers, 5x First-strand Buffer, 0.1 M DTT, 40 units of RNase OUT and 200 units of M-MLV RT, in a volume of 20 μL for each reaction (all Invitrogen). qPCR (Bio-Rad) was performed using Bio-Rad iQ SYBR Green Supermix according to manufacturer’s instructions. For each sample, PCR buffer was mixed with 100 ng cDNA, sybergreen and forward and reverse primers for the targeted genes in a volume of 13 μL. A three-step qPCR reaction was applied. Oligo sequences of primes used were as followed: CALCA fwd, 5′-GGCTTCCAAAAGTTCTCCCCC-3′; CALCA rev, 5′-CCAGCCGATGAGTCACACAG -3′; CEACAM5 fwd, 5′-AAGAAATGACGCAAGAGCCTATG-3′; CEACAM5 rev, 5′-CCCGAAAGGTAAGACGAGTCTG-3′; SST fwd, 5′-ACCCAACCAGACGGAGAATGA-3′; SST rev, 5′-GCCGGGTTTGAGTTAGCAGA-3′, TTF1 fwd, [ATGTACCGGGACGACTTGGAA]; TTF1 rev, [CAATGCCTGTCAGGGCTAGAA] and YWHAZ fwd, 5′-GATCCCCAATGCTTCACAAG-3′; YWHAZ rev, 5′-TGCTTGTTGTGACTGATCGAC -3′.

### Immunostaining

For immunostaining, antibodies to Calcitonin (1:100, AB_2335678), CEA (1:100, AB_2244691), TTF-1 (1:100, AB_1310784), and SST (1:100, AB_2195910) were used to detect proteins. In the case of the organoids, Matrigel was dissolved by incubation with Dispase enzyme. Followed by subsequent washes with PBS/0.2% BSA and centrifuged at 400 g for 5 min. The resulting pellet was fixed in 4% paraformaldehyde (15 min, RT) and washed with PBS. Next, the organoids were embedded in HistoGel (Richard-Allan Scientific/Thermo scientific) and the gel was subjected to dehydration, followed by embedding in paraffin and sectioning (5 μm). Patient material was fixed in 4% formaldehyde. After dehydration, the tissue was paraffin-embedded and sectioned at a 5 μm thickness. Sections were de-paraffinized and subsequently, Tris-EDTA antigen-retrieval was performed (pH 9.0). This was followed by washing, blocking, and incubation with primary antibodies overnight. Slides were then incubated with secondary antibodies, stained with 4′,6-diamidino-2-phenylindole (DAPI), and mounted with either aqueous mounting medium (DAKO) for tissue sections or prolong glass antifade mountant (ThermoFisher) for organoid sections.

### Microscope image acquisition

Fluorescent microscopy was performed at room temperature with a white light laser confocal microscope (Leica TCS SP8 X) equipped with a sCMOS camera (Leica DFC9000 GTC). Images were acquired with a 63x oil objective (Leica HC PL APO CS2, NA:1.4) using Las X software. Imaging was performed using stained sections mounted using aqueous mounting medium (DAKO) for tissue sections or prolong glass antifade mountant (ThermoFisher) for organoid sections. As fluorochromes the following were used: Goat anti-Rabbit IgG (H+L) Cross-Adsorbed Secondary Antibody, Alexa Fluor™ 633 (Invitrogen), and Goat anti-Mouse IgG (H+L), Superclonal™ Recombinant Secondary Antibody, Alexa Fluor™ 633 (Invitrogen). The images were analyzed by ImageJ (National Institutes of Health).

### Tyrosine kinase inhibitor screening

Vandetanib (Selleck Chemicals), cabozantinib (LKT Laboratories Inc.), selpercatinib (Selleck Chemicals) and pralsetinib (Selleck Chemicals) were added to the medium of the MZ-CRC-1 MTC cell-line derived spheroids, cultured in an Ultra-low attachment U-bottom plate, in a logarithmic dose de-escalation per TKI, one week after start of culture. Incucyte images during 72 hours after TKI administration were obtained and used to analyze cytotoxicity induced by the TKI. The Largest Brightfield Object Area (um^2^) metric was used to determine the IC50, the final concentration per TKI was determined and used for further TKI screening in patient-derived MTC organoids (**Supplemental Figure 1**). The final dose of 1.0 μM of vandetanib and 0.1 μM of cabozantinib, selpercatinib, and pralsetinib were added to the medium of MTC organoids at the end of passage 1. After 72 hours, medium was collected to measure concentration of secreted calcitonin and CEA.

### PET tracer validation

To demonstrate the utility of patient-derived MTC organoids to validate PET/SPECT tracers, existing tracers, i.e., ^18^F-fluorodeoxyglucose (^18^F-FDG), ^18^F-L-3,4-dihydroxyphenylalanine (^18^F-DOPA), and ^18^F -Prostate Specific Membrane Antigen (^18^F-PSMA) were used to assess uptake of the tracer by the MTC organoids, specific uptake was determined by using a 5% glucose solution, a cold dose of 0.76 mM DOPA, and a cold dose of 0.38 mM PSMA. Prior to the experiments, the Matrigel was dissolved by incubation with Dispase enzyme followed by three washes with 1000 μL of HBSS/1% BSA. Approximately 10,000 organoids in 450 μL HBSS/1% BSA were incubated with 1 MBq (in 50 μL) tracer solution at 37 °C for 60 minutes. After tracer incubation, the organoids were collected with a 5-micron filter needle and subsequently washed three times through the filter needle with 1000 μL ice-cold HBSS/1% BSA. Activity in the organoids was measured in a gamma-counter (Wizard^2^ 2480, SW 2.1, PerkinElmer). Tracer uptake was corrected for the number of organoids by counting the number of organoids using the Ocular Micrometer at a 10x objective prior to the experiment. Uptake was expressed as percentage of organoid-associated radioactivity per 10,000 organoids. Uptake values were corrected for background radiation. All experiments were executed in triplicate.

### Statistics

All data represent the mean ± SEM. Statistical analyses and producing graphs were done using GraphPad Prism, version 8 (GraphPad Software). The statistical methods relevant to each figure are described in the figure legends. Statistical comparisons between two groups were performed using the Mann-Whitney U test or Student’s t test (depending on normal distribution). P values of less than 0.05 were considered to indicate a significant difference between groups.

### Study approval

Written informed consent of participants was obtained prior to participation. The study was approved by the Medical Ethics Committee Groningen (approval no. 2015/101).

## Author contributions

Conceptualization, L.H.J.S., E.C.J., R.P.C., and S.K.; methodology, L.H.J.S., and E.C.J. ; formal analysis, L.H.J.S., E.C.J., I.F.A., R.M.; investigation, L.H.J.S., E.C.J., I.F.A. R.M.; resources, I.F.A., L.J., W.T.Z., A.H.B., T.P.L., R.P.C., and S.K.; data curation, L.H.J.S., E.C.J., R.P.C., and S.K.; writing – original draft, L.H.J.S., and E.C.J.; writing – review & editing, L.H.J.S., E.C.J., I.F.A., R.M., L.J., W.T.Z., A.H.B., T.P.L., R.P.C., and S.K.; visualization, L.H.J.S., and E.C.J.; supervision, R.P.C., A.H.B., T.P.L., and S.K.; project administration, R.P.C. and S.K.; funding acquisition, T.P.L., R.P.C., and S.K.

## Acknowledgements

This work was partly funded by UMCG Cancer Research Fund 490178 (to R.P.C. and S.K.) and 4901250 (T.P.L.) and by Stichting De Cock-Hadders 2023-60 (to R.P.C., T.P.L., and S.K). Part of the work was performed at the UMCG Imaging and Microscopy Center (UMIC).

## Conflict of interest

The authors declare no competing interests.

## Notes

### Competing Interest Statement

The authors have declared no competing interest.

