## Supplemental figures for "Patient-Derived Medullary Thyroid Cancer Organoids; a Model for Patient-tailored Drug and PET-Tracer Screening"

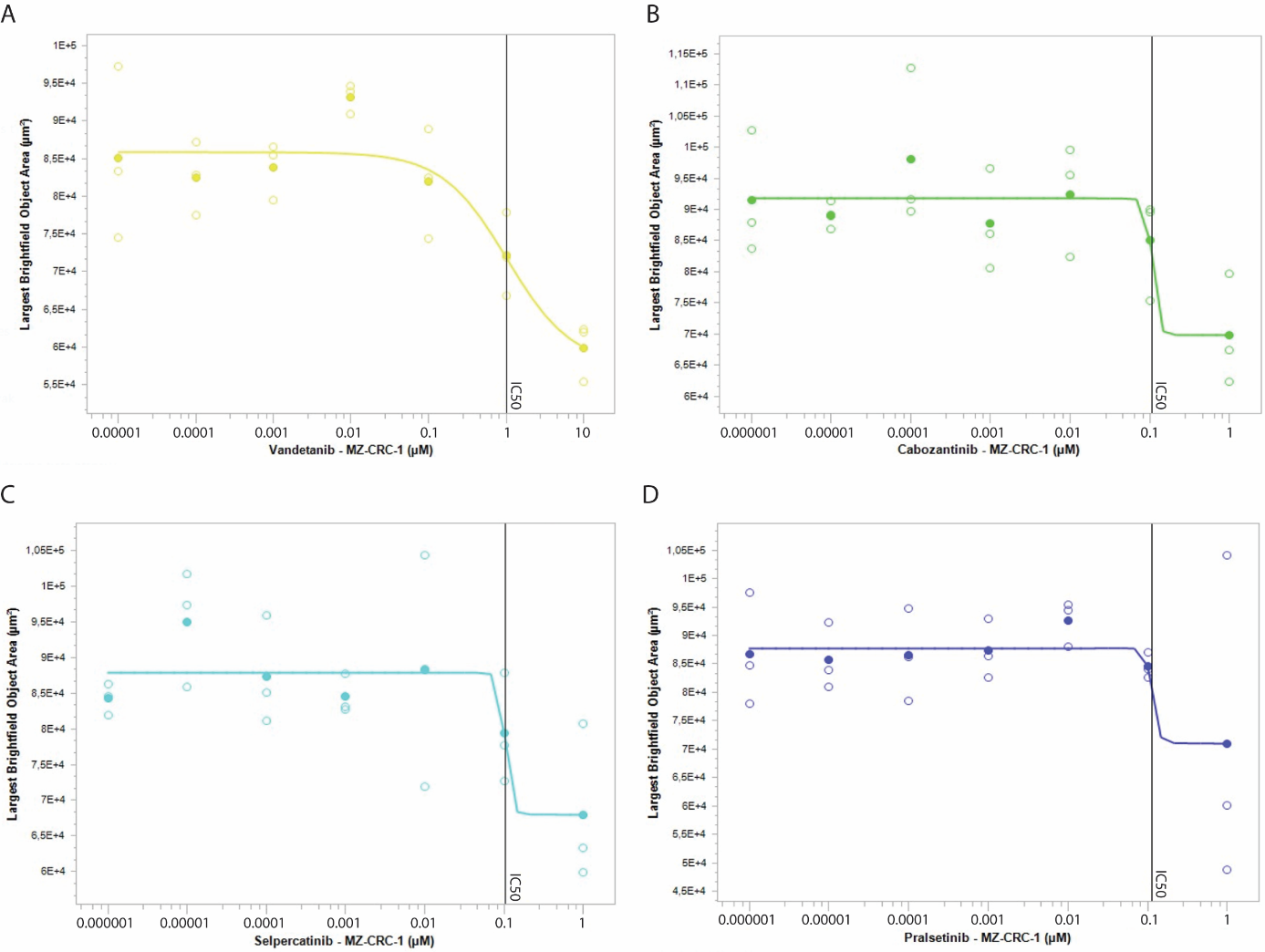

*Supplemental Figure 1. Determination of the IC50 value of (A) Vandetanib; 1µM, (B) cabozantinib; 0.1 µM, (C) selpercatinib; 0.1 µM, and (D) pralsetinib; 0.1 µM, on the 3D-cultured MTC cell line MZ-CRC-1. Spheroids were cultured in an Ultra-low attachment U-bottom plate, in a logarithmic dose de-escalation per TKI. Images were obtained during 72 hours of exposure, and used to analyze cytotoxicity induced by the TKI. The Largest Brightfield Object Area (um2) metric was used to determine the IC50, the final concentration per TKI was determined and used for further TKI screening in patient-derived MTC organoids.*
